# CNV-PG: a machine-learning framework for accurate copy number variation predicting and genotyping

**DOI:** 10.1101/2020.04.13.039016

**Authors:** Taifu Wang, Jinghua Sun, Xiuqing Zhang, Wen-Jing Wang, Qing Zhou

## Abstract

**Motivation:** Copy-number variants (CNVs) are one of the major causes of genetic disorders. However, current methods for CNV calling have high false-positive rates and low concordance, and a few of them can accurately genotype CNVs.

**Results:** Here we propose CNV-PG (CNV Predicting and Genotyping), a machine-learning framework for accurately predicting and genotyping CNVs from paired-end sequencing data. CNV-PG can efficiently remove false positive CNVs from existing CNV discovery algorithms, and integrate CNVs from multiple CNV callers into a unified call set with high genotyping accuracy.

**Availability:** CNV-PG is available at https://github.com/wonderful1/CNV-PG

## 1 Introduction

Copy-number variants (CNVs) are one of the major causes of genetic disorders^[1]^, making accurate detection of CNV essential for diagnosis of such diseases. Currently, many next-generation sequencing (NGS)-based CNV detection methods have been proposed^[2]^. However, most of these show high false-positive rates because of the noises in the sequencing data, such as sequencing error and artificial chimeric reads, and ambiguous mapping of reads cause by repeat- and duplication-rich regions^[2]^. To identify a set of high-confidence CNVs, a strategy that takes intersecting CNVs generated by two or more algorithms is widely used. However, due to different CNV-property-dependent and library-property-dependent features use by CNV detection methods, they show low concordance, causing a large number of potentially true CNVs to be discarded. Besides, a few of present software can accurately give the genotype of a CNV, causing a challenge for accurate detection of de novo CNVs. Here, we present CNV-PG, a machine-learning framework that aims at accurately predicting and genotyping true CNVs from identified results by various software. CNV-PG an open-source application written in Python, including two parts **(Figure 1)**: CNV predicting (CNV-P) and CNV genotyping (CNV-G). For CNV-P, we trained a model on a subset of validated CNVs from 5 commonly used software for CNV detection separately, and obtained the corresponding classifier for predicting true CNVs. For CNV-G, providing accurate genotypes for CNVs, it is compatible with existing CNV detection algorithms.

**Figure 1:**
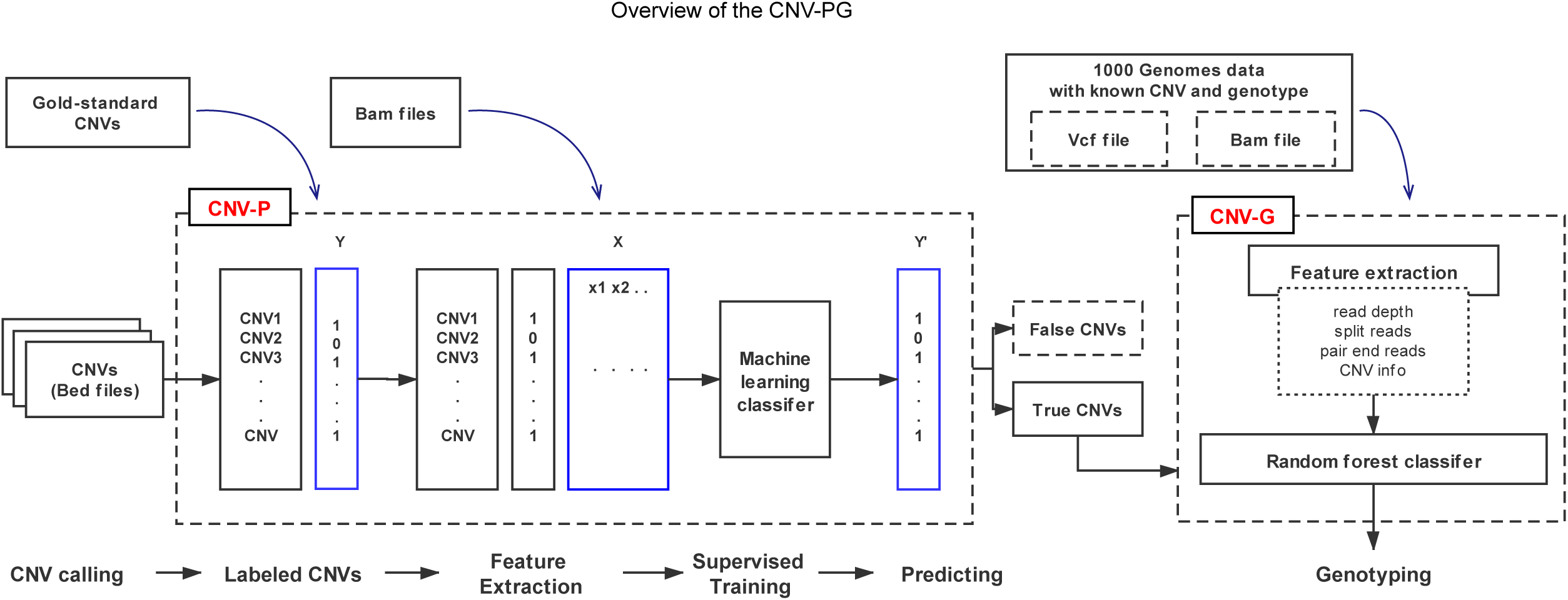
Overview of the CNV-PG. The CNV-PG consists of two parts: CNV predicting (CNV-P) and CNV genotyping (CNV-G). In CNV-P, we trained a supervised machine learning model to classify candidate CNVs as True or False. Then, CNV-G was performed to accurately give the genotypes of these high-confidence CNVs.

## 2 Methods

### 2.1 Data sets

In CNV-P, The gold-standard CNV sets of 9 individuals (NA19238, NA19239, NA19240, HG00512, HG00513, HG00514, HG00731, HG00732, HG00733) were download from Chaisson et al 2019^[3]^. The whole genome sequences (WGS) data (∼30x) of these 9 individuals were downloaded from National Center for Biotechnology Information (NCBI) with accession number of SRP159517 (**Supplemental table. S1, S2**). For Validation sets, the sequencing data of NA12878 and HG002 were also downloaded from NCBI with accession number SRP159517 and SRP047086 respectively. The gold-standard CNV dataset for NA12878 was generated by three data sets: the Database of Genomic Variants (http://dgv.tcag.ca/dgv/app/home?ref=GRCh37/hg19)^[4]^, the 1000 Genomes Project phaseIII (https://ftp.ncbi.nih.gov/1000genomes/ftp/phase3/integrated_sv_map/) ^[5]^, and the PacBio CNV data from Pendleton, M. et al.2015^[6]^. The gold-standard CNV dataset for HG002 was downloaded from Zook, J. M. et al.2019^[7]^.

In CNV-G, validated genotypes and aligned bam files for 26 individuals were downloaded from the 1000 Genomes Project (https://ftp.ncbi.nih.gov/1000genomes/ftp/phase3/data/). Sequencing data of validation sample NA12878 was downloaded from NCBI with accession number SRR7782683 and its genotypes were collected from Conrad. D. F. et al.2009^[8]^ (**Supplemental table. S1, S3**).

### 2.2 Predicting

A total of five commonly-used softwares (Lumpy^[8]^, Manta^[9]^, Pindel^[10]^, Delly^[11]^ and breakdancer^[12]^) were chosen in our study. Although these software detect CNVs based on different variables, such as Pindel using the signal of split reads and Breakdancer using the information of paired reads, there are consistent features for a certain CNV. We choose commonly used features in all software to characterize CNVs in our training model, including size, read depth, information of paired and spited read, and GC content of CNV body, as well as all these features around CNV’s boundaries (**Supplemental Table S1, S2**).

### 2.3 Genotyping

Training features were collected based on seven informative signals: depth of coverage, GC content, split-reads, discordant paired-ends, CNV size ranges, CNV type, and the number of chromosomes. All of these characteristics may contribute to genotyping (**Supplemental Table S1, S3**).

## 3 Results

### 3.1 CNV-P identified a set of high-confidence CNVs with high precision and recall rates

To illustrate the efficiency and characteristics of the CNV-P, we used CNVs from six samples in Chaisson et al 2019^[3]^ for training, the remaining 3 samples for evaluation **(Supplemental Table. S2)**. We first identified CNVs of nine samples using five frequently-used CNV callers (Lumpy^[8]^, Manta^[9]^, Pindel^[10]^, Delly^[11]^ and breakdancer^[12]^) respectively. For each CNV set, we removed CNVs with low quality and locating on N region of genome to get the “row CNVs”. Then, we labeled CNVs as either “True” or “False” based on a 50% reciprocal overlap with the gold-standard CNVs. Finally, the labeled CNVs were used to train CNV-P using a Random Forest classifier and the remaining CNVs were used to evaluate its performance **(Fig. 1)**. we trained the CNV-P classifier on 10-fold cross-validation for optimal parameter selection (**Supplemental Fig. S1**). Thus, we obtained a Random Forest classifier for each CNV detection method.

Using the evaluation set mentioned above, each caller-specific CNV-P classifier realized accurately classified the CNVs as either true or false at over 91% precision (95% for Lumpy, 93% for Manta, 93% for Pindel, 92% for breakdancer, 91% for Delly) and over 87% recall rates (96% for Lumpy, 95% for Manta, 93% for Pindel, 95% for breakdancer, 87% for Delly). The overall diagnostic ability of each classifier, measured as the area under the Receiver Operating Characteristic (ROC) curve (AUC), was 97% for Lumpy, 94% for Manta, 97% for Pindel, 93% for breakdancer, and 96% for Delly (**Supplemental Fig. S2.A, B**). Additionally, we noticed that after our classification, a large number of false positive CNVs were removed, and majority of the true CNVs were remained (**Supplemental Fig. S2.C**). To dissect the principle of the CNV-P classifier, we assessed the relative importance of each feature for corresponding classifiers. As expected, for all classifiers, read-depth provided the most discriminatory power to make accurate CNV predictions (**Supplemental Fig. S3**). While the second important feature inconsistent in different classifiers, it may reflect caller-specific CNV signals.

To evaluate the robustness of each CNV-P, we trained each CNV-P on varying proportions of training data (from 10% to 90% in increments of 20%). The results show a steady improvement in accuracy (precise and recall rate) with an increase in the number of training data (**Supplemental Fig. S4**). CNV-P performed well based on even 10% of training sets, showing over 90% precise rate and 87% recall rate. We further assessed the performance of CNV-P for different size of CNVs. We divided CNVs into three sets based on their size: CNV_S (100bp to 1kb), CNV_M (1kb to 100kb), CNV_L (>100kb). The overall precision of each size interval was greatly improved, comparing with the row CNVs achieved by the corresponding CNV callers (**Supplemental Fig. S5**). We noticed that almost all precise and recall rate in the size range of CNV_S and CNV_M were over 90%, while the theses value in CNV_L was slight lower. This may be due to the insufficient number of CNV_L in our training data, since they all come from healthy individuals who do not have a lot of large size of CNVs. As a result, each input CNV would get a probability score predicted by CNV-P, it can be used as a measurement of CNV confidence (**Supplemental Fig. S6**).

We also implanted two additional predictors to CNV-P, Gradient Boosting classifier (GBC) and Support Vector Machine (SVM) classifier. When compared these different supervisor machine learning classifiers, we found little qualitative difference between GBC and Random Forest Classifier, Random Forest showed slightly better performance, and SVM was the worst performer (**Supplemental Fig. S7**).

To further validate the performance of CNV-P, we implemented two independent WGS datasets from NA12878 and HG002 (**Supplemental Table. S1**). In this part, each caller-specific classifier was trained on data of all nine individuals mentioned above. Consistent with the above results, CNV-P produced the optimal performance with AUCs of 0.94,0.93,0.93,0.88 and 0.95 for Lumpy, Manta, Delly, Pindel and breakdancer respectively in NA12878 (**Fig. 2A**). Most of false-positive CNVs were removed with a small true positive loss **(Fig. 2B, C)**. Likewise, HG002 presents the same performance (**Fig. 2E-G**). Moreover, CNV-P also showed a good performance on sequencing data generate by BGI-500 sequencing platform **(Supplemental Fig. S8)**.

**Figure 2:**
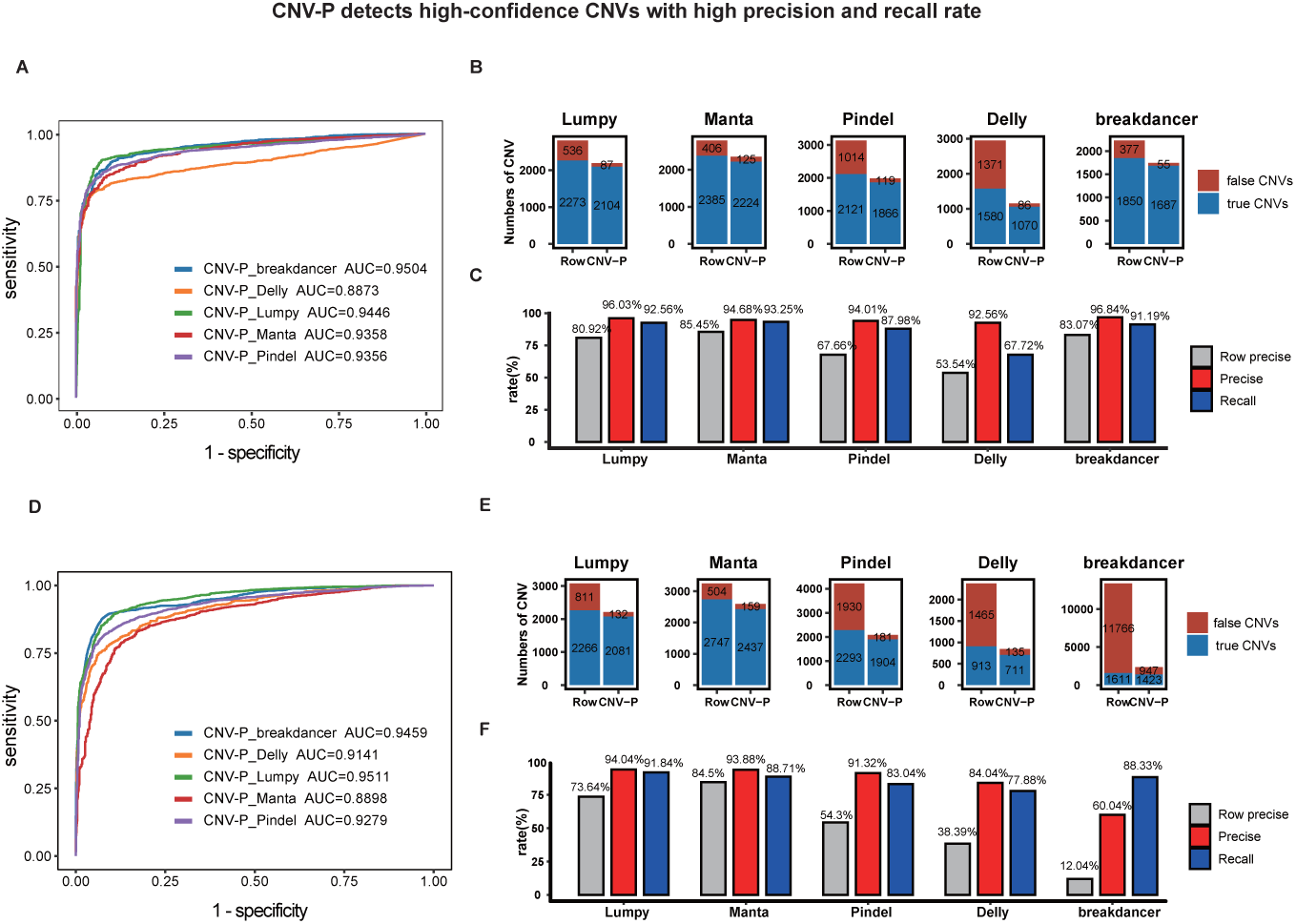
CNV-P detects high-confidence CNVs with high precision and recall rates. A, D) Receiver operating characteristic (ROC) curves of CNV-P; B, E) The number of classified CNVs by CNV-P from five commonly used tools; C, F) The precise and recall of CNV-P;

### 3.2 CNV-G provide accurate genotypes of CNVs

Many CNV callers have a function of genotyping CNVs, such as Manta, Delly and Lumpy with a companion tool svtyper^[13]^. However, some other software did not provide genotypes for CNVs. Here, we also developed a machine-learning approach, named CNV-G, for genotyping a certain CNV. Model selection was performed on training data using 10-fold cross-validation (**Supplemental Fig. S9**). The classifier performance was independently evaluated in the NA12878.

To verify the performance of CNV-G, we compared it to several widely used CNV genotyping tools including svtyper, Manta and Delly. We use Delly, Manta and Lumpy&svtyper to generate an initial CNV set. Then, CNV-G genotyped the union from Delly, Lumpy and Manta for the validation set of NA12878 whose genotypes generated by the Agilent 105K CNV genotyping array. We generated a receiver operating characteristic (ROC) curves for each genotyping method. Also, we implanted 3 predictors to CNV-G, Random Forest Classifier (RF), Gradient Boosting classifier (GBC) and Support Vector Machine (SVM) classifier. CNV-G-RF (CNV-G based on Random Forest) produced the best genotyping accuracy with AUCs of 0.95, in contrast to CNV-G-GBC (CNV-G based on Gradient Boosting) and CNV-G-SVM (CNV-G based on Support Vector Machine) (**Fig. 3**). Likewise, when compared to other genotyping methods, we found that CNV-P-RF produced the optimal performance with AUCs of 0.95, while Manta performance resulted in an AUC of 0.91, Delly and svtyper producing AUCs of 0.93 and 0.82, respectively.

**Figure 3:**
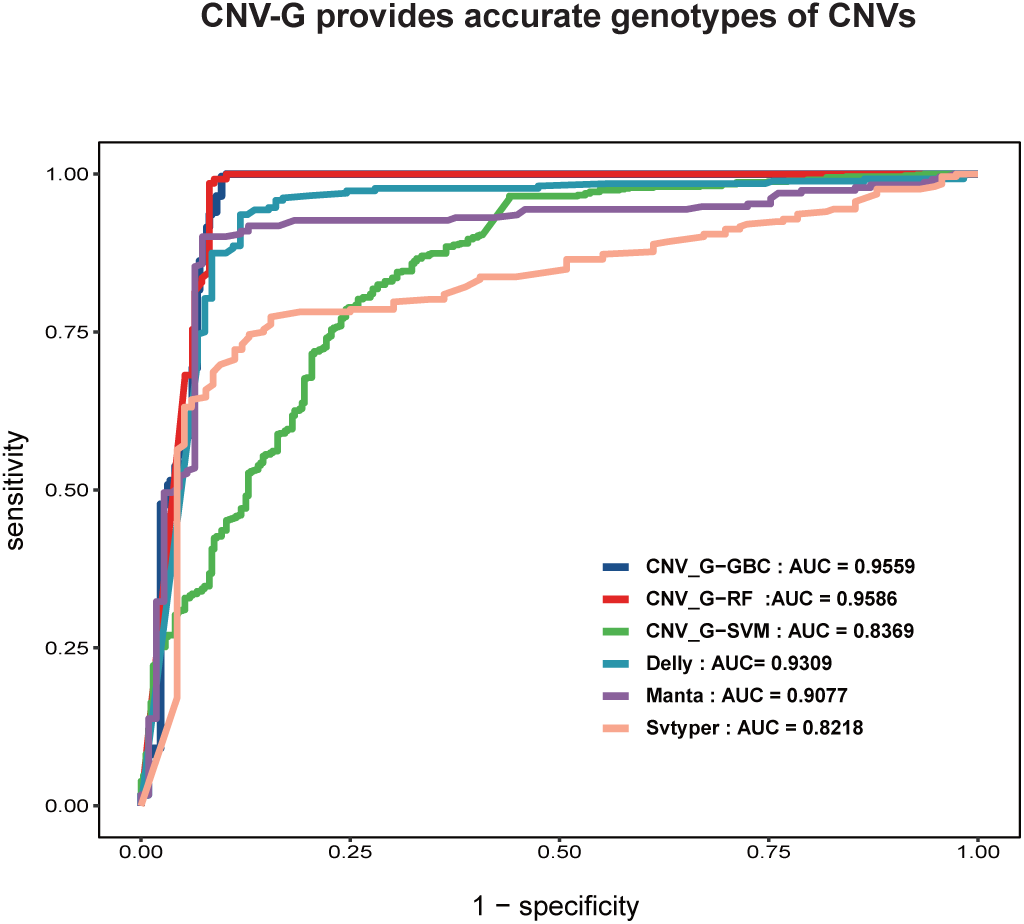
CNV-G provides accurate genotypes of CNVs. Receiver operating characteristic (ROC) curves of genotyping for CNV-G (RF-based, GBC-based, and SVM-based), Delly, Svtyper, and Manta;

## Conclusions

CNV detection from WGS is error-prone because of short-read length and library-property-dependent bias. Inflated false positives making a big challenge for researchers to identify clinically relevant CNVs, as it is time and money consuming to validate a large amount of false positive CNVs. To solve this problem, we provide CNV-PG, an effective machine-learning-based framework to acquire high-quality CNVs and their genotypes. Instead of handling the shortcomings of existing methods by developing another CNV caller, CNV-PG focused on creating a reliable integrative CNV set from existing CNV detection software. We demonstrate that CNV-PG can identify a set of high-confidence CNVs with high precision and recall rates, and the accuracy of genotypes outperform present widely used CNV genotyping tools. Moreover, CNV-PG is robust to variation in the proportion of training sets, CNV size and sequencing platforms, indicating the utility of CNV-PG in a variety of clinical or research contexts.

The limitations of CNV-PG, is its dependency on a set of validated CNVs from several healthy individuals. Therefore, there were not enough large size CNVs in our training data and may have weaker power for large-size CNVs. Even though, our results demonstrate the utility of CNV-PG, which can serve as a proof-of-principle for future studies that accumulate enough large pieces of ‘gold standard’ CNVs curated from some disease samples as training data.

Overall, CNV-PG provides a well-performed machine-learning framework for accurately predicting and genotyping CNVs, which make great sense to generate a set of high-confidence CNVs, and benefit both the basic research and clinical diagnostic of genetic diseases.

## Acknowledgements

The authors thank Dr. Jian Guo for constructive comments on this project and Chen Ye for data download and management.

## References

1. Stankiewicz, P. and J.R. Lupski, Structural variation in the human genome and its role in disease. Annu Rev Med, 2010. 61: p. 437–55.

2. Kosugi, S., et al., Comprehensive evaluation of structural variation detection algorithms for whole genome sequencing. Genome Biol, 2019. 20(1): p. 117.

3. Multi-platform discovery of haplotype-resolved structural variation in human genomes. Nature Communications, 2019.

4. R., M.J., et al., The Database of Genomic Variants: a curated collection of structural variation in the human genome. Nucleic Acids Research, 2013(D1): p. D1.

5. An integrated map of structural variation in 2,504 human genomes. Nature. 526(7571): p. 75–81.

6. Pendleton, M., et al., Assembly and diploid architecture of an individual human genome via single-molecule technologies. Nature Methods. 12(8): p. 780–786.

7. Zook, J.M., et al., A robust benchmark for germline structural variant detection. bioRxiv, 2019: p. 664623.

8. Layer, R.M., et al., LUMPY: a probabilistic framework for structural variant discovery. Genome Biology. 15(6): p. R84.

9. Chen, X., et al., Manta: Rapid detection of structural variants and indels for germline and cancer sequencing applications. Bioinformatics, 2015. 32(8): p. 1220–1222.

10. Ye, K., et al., Pindel: a pattern growth approach to detect break points of large deletions and medium sized insertions from paired-end short reads. Bioinformatics. 25(21): p. 2865–2871.

11. Rausch, T., et al., DELLY: structural variant discovery by integrated paired-end and split-read analysis. Bioinformatics, 2012. 28(18): p. i333.

12. Chen, K., et al., BreakDancer: an algorithm for high-resolution mapping of genomic structural variation. Nat Methods, 2009. 6(9): p. 677–81.

13. Chiang, C., et al., SpeedSeq: ultra-fast personal genome analysis and interpretation. Nat Methods, 2015. 12(10): p. 966–8.

