## Supplemental information for "CNV-PG: a machine-learning framework for accurate copy number variation predicting and genotyping"

---

### *Supplementary Method*

### 1. CNV Calling processes

For the WGS data, we aligned its to the hg19 reference with bwa(bwa-0.7.16a) 'mem' command to generate the alignment BAM file. Then, the BAM file was submited to CNV callers as input data. The commands and options used for the 5 CNV callers used in this study are described below(Taking HG002 for example).

#### 1.1 Lumpy

For running Lumpy, we implanted SpeedSeq workflow, we firstly extracted discordant paired read alignments and split read alignments with samtools and the extractSplitReads_BwaMem command. The 'lumpy' command was executed with the options (eg. HG002):

-pe

bam_file:HG002.discordants.bam,histo_file:HG002.lib1.x4.histo,mean:569.0479672525416,stdev:156.96862863277676,read_length:148,min_non_overlap:148,discordant_z:5,back_distance:10,weight:1,id:HG002,min_mapping_threshold:20,read_group:HG002

-sr

bam_file:HG002.splitters.bam,back_distance:10,min_mapping_threshold:20,weight:1,id:HG002,min_clip:20.

#### 1.2 manta

For running Manta, we run 'configManta.py' script on the input bam file and hg19 reference file, then, the 'runWorkflow.py' script was run to execute the entire workflow. The 'manta' was executed with the commands:

1) $Bin/configManta.py --bam HG002.bam --referenceFasta hg19.fa --runDir ./

2) $Bin/runWorkflow.py -j 8

#### 1.3 delly

For running delly, we run 'delly call' command on the input bam file and hg19 reference file to get the BCF file, then bcftools was executed for converting BCF to VCF.

commands used:

1) $Bin/delly call -t ALL -g hg19.fa -o HG002.delly.bcf HG002.bam
2) $Bin/bcftools view HG002.delly.bcf > HG002.delly.vcf

#### 1.4 pindel

For running Pindel, we created a configure file for each chromosome, in which the input bam file and the mean insert size were specified. Then, the 'pindel' command was executed for each chromosome with the hg19 reference file and configure file. Finally, pindel2vcf was executed for converting output files to a VCF file.

commands used(eg. chromosome 1):

1) $Bin/pindel -f hg19.fa -i HG002. Configure_file -c chr1 -T 16 -x 2 -M 3 -v 100 -d 30 -E 0.92 -w 10 -o HG002.chr1
2) $Bin/pindel2vcf -r hg19.fa -R hg19 -P HG002.chr1 -v HG002.chr1.sv.vcf

#### 1.5 Breakdancer

For running breakdancer, we executed 'bam2cfg.pl' script to create configure files. Then, breakdancer-max was executed for all chromosomes.

commands used:

1) $Bin/ bam2cfg.pl -v 20 HG002.bam > HG002.SV.cfg
2) $Bin/ breakdancer_max -y 30 -x 1000 -r 2 -m 10000000 HG002.SV.cfg > HG002.SV.ctx

### 2. Training sets and validation sets

In CNV-P, we firstly filter and merge the CNVs for each CNV caller with the below criteria: 1) Removing CNVs overlapped(> 10bp) with N region (download from UCSC: http://genome.ucsc.edu/) 2) if two CNVs exhibited ≧ 80% reciprocal overlaps, we merge them to one CNV, and the smaller start and larger end to be the new start and end of the merged CNV. 3) Removing CNVs that length less than 100bp. Then, we measured the extent of overlap among callers for each CNV event and labeled the calls as either 'True' or 'False' based on their overlap with gold-standard CNV calls. The 'True' CNV calls were judged when the called CNVs exhibited ≧ 50% reciprocal overlaps with the gold-standard CNVs. Training features were obtained from the BAM files, and the details are shown in **Supplementary Table S3**.

In CNV-G, the training sets were obtained from 26 PCR-free high coverage whole genomes and the 1KGP phase 3 CNV genotypes as truth. The training features are shown in **Supplementary Table S4.** For evaluating the performance of CNV-G, Delly, Lumpy combined with svtyper, and Manta were used to generate an initial CNV call set. Then, CNV-G genotyped the union of Delly, Lumpy and Manta calls for the validation set of NA12878(The genotypes of its generated by the Agilent 105K CNV genotyping array). The 'True' genotype calls were judged when the called CNVs exhibited ≧ 80% reciprocal overlaps with the gold-standard CNVs and share with the same genotypes.

### *Supplementary Figures*

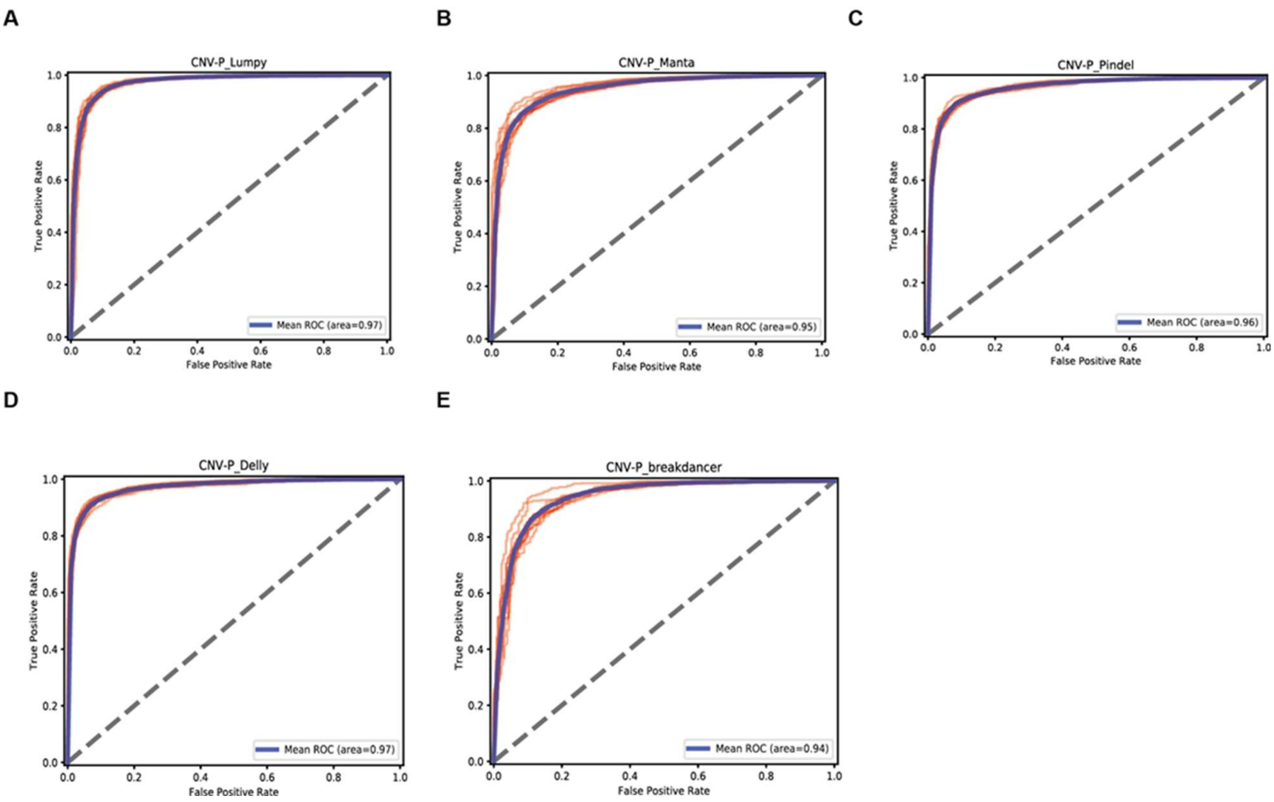


**Figure S1. 10-fold Cross validation for CNV-P.** For each classifier, we performed cross-validation of the training set in ten folds. A) Lumpy; B) Manta; C) Pindel; D) Delly; E) breakdancer. The red line showed the ROC for each cross-validation and blue lines showed the mean ROC curve.

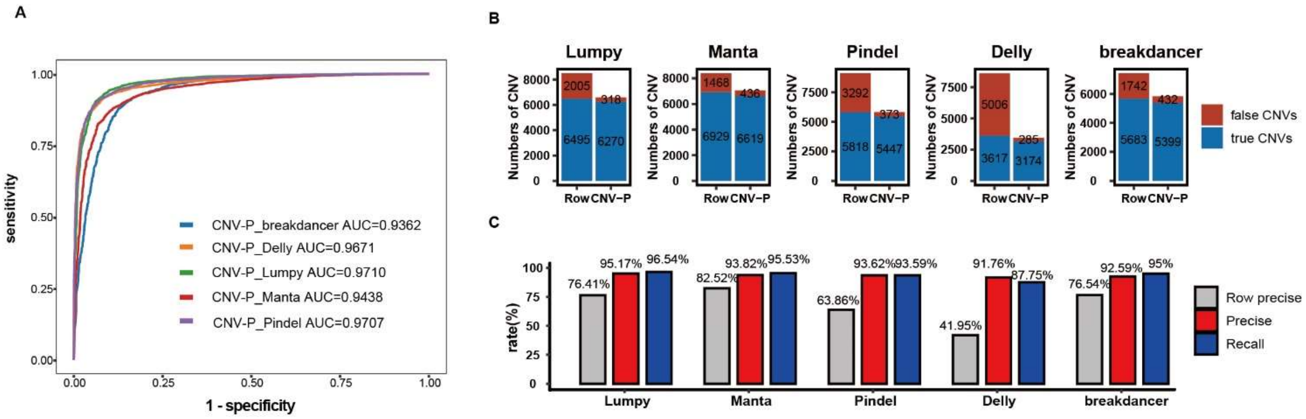


**Figure S2. Performance of CNV-P in 3 test samples.** A) Receiver operating characteristic (ROC) curves of CNV-P in 3 test samples. B)The number of CNVs before or after CNV-P predicting for five commonly used tools. C) The precise and recall of CNV-P.

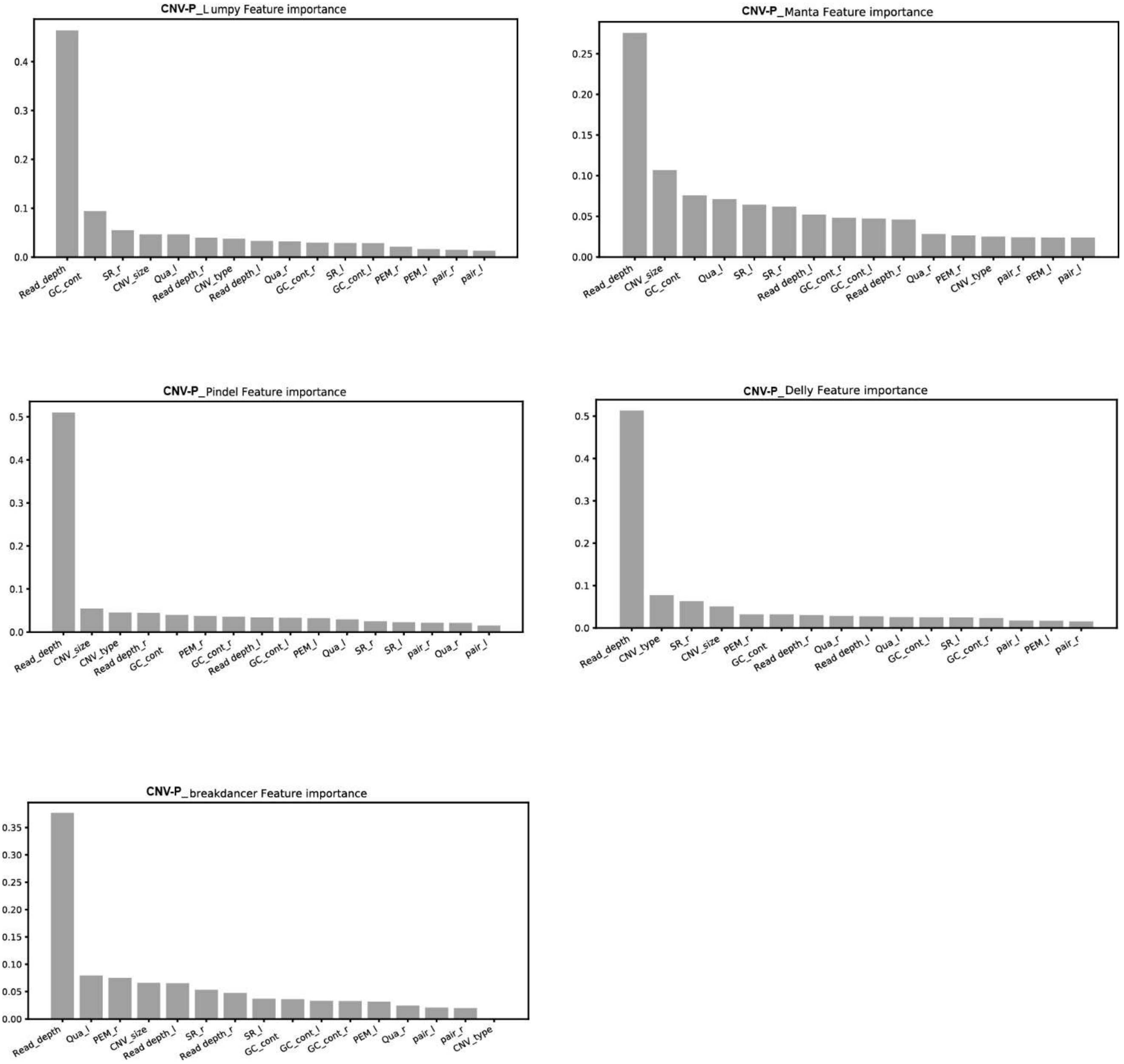


**Figure S3. CNV-P Features importance. We train a CNV-P** classifier for each CNV caller base on Random Forest, the relative importance of each features supplemented to CNV-P is shown.

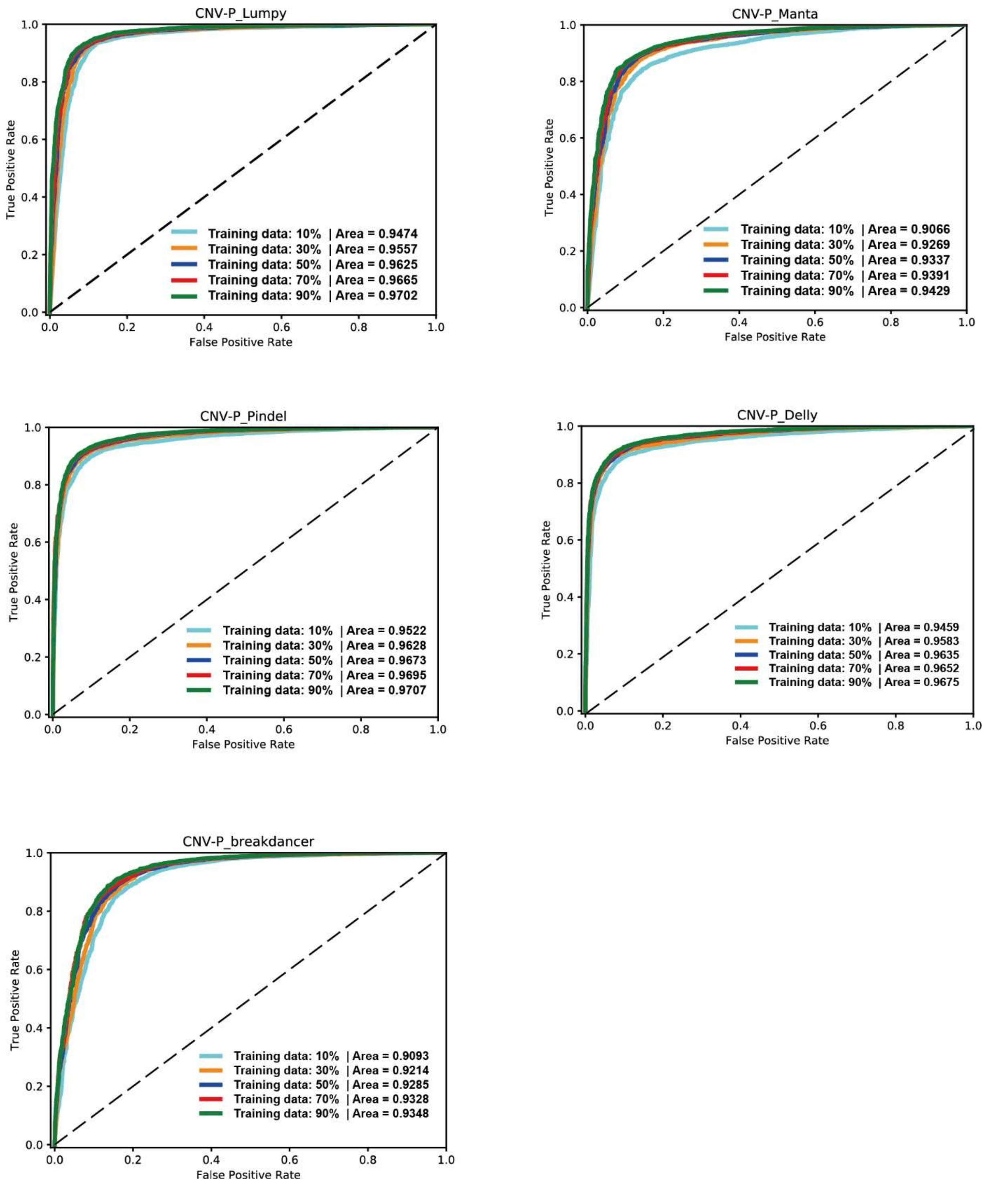


**Figure S4. Classifier improved with increase of training data.** To evaluate the robustness of each CNV-P, we trained each CNV-P on varying proportions of training data (from 10% to 90% in increments of 20%). the ROC of each CNV-P classifier is shown.

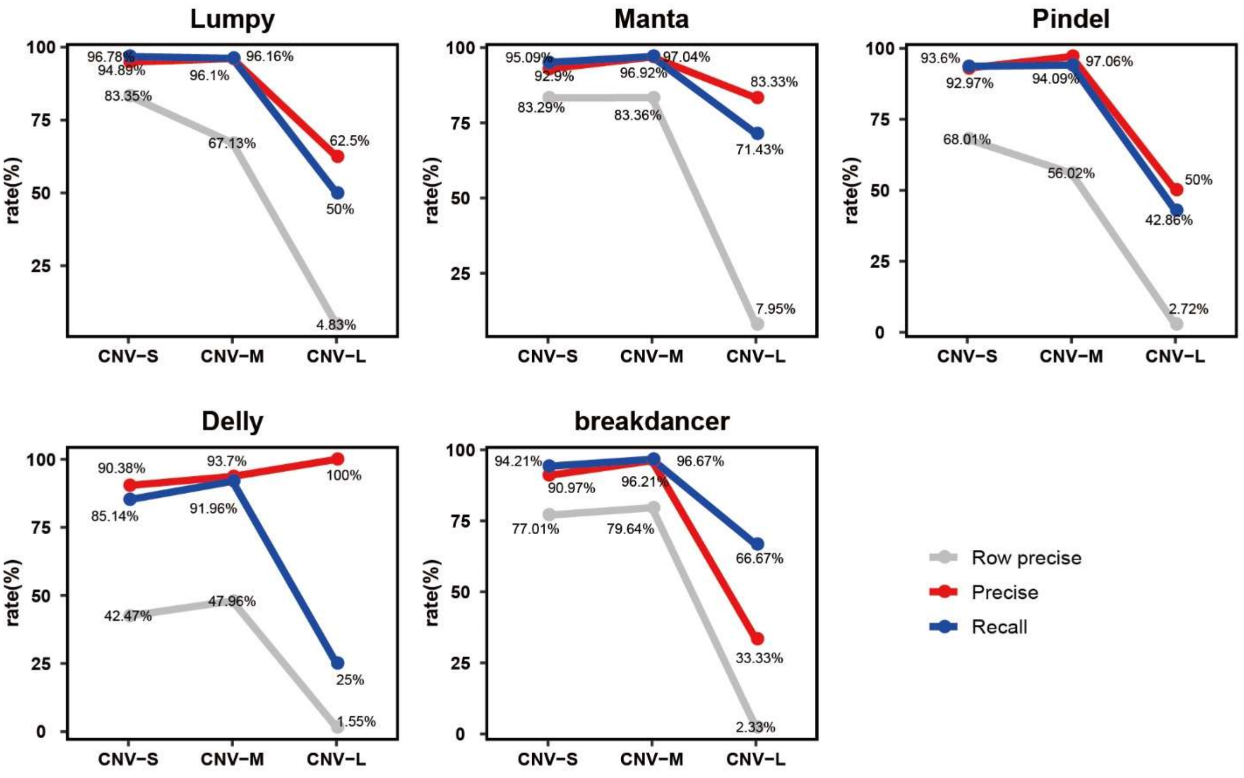


**Figure S5. Performance of CNV-P at different size ranges.** We divided CNVs into three sets based on their size: CNV_S (100bp to 1kb), CNV_M (1kb to 100kb) and CNV_L (>100kb). For each CNV-P, precise and recall rate in the size range of CNV_S and CNV_M were over 90%, while the theses value in CNV_L was slight lower.

A) CNV-P_Lumpy

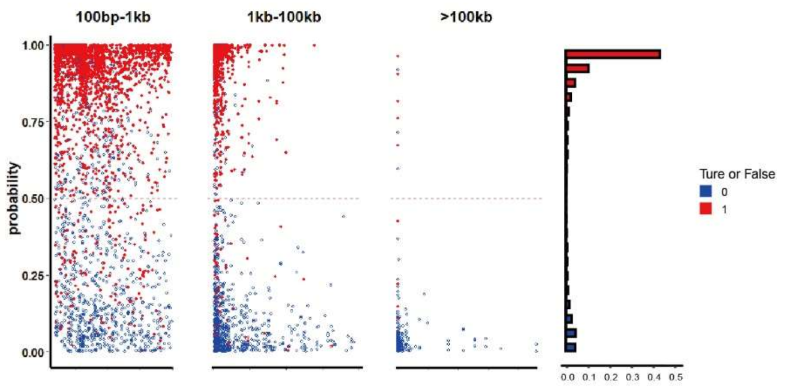


B) CNV-P_Manta

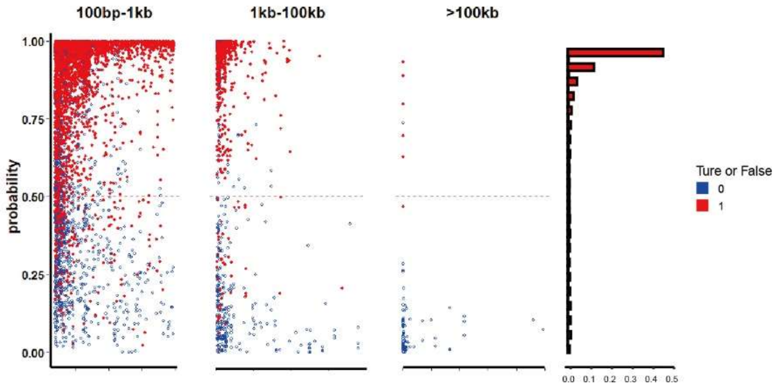


C) CNV-P_Pindel

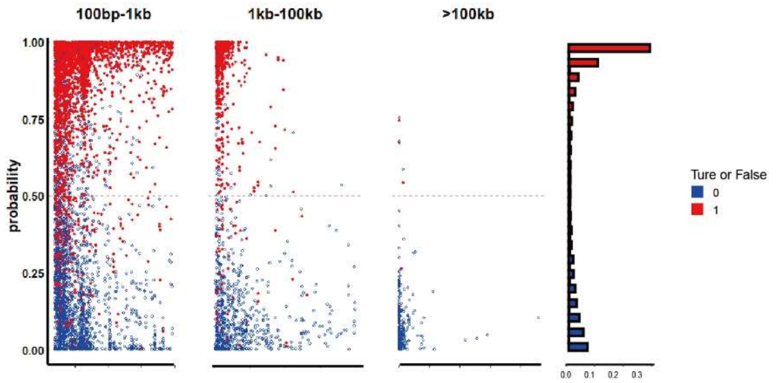


D) CNV-P_Delly

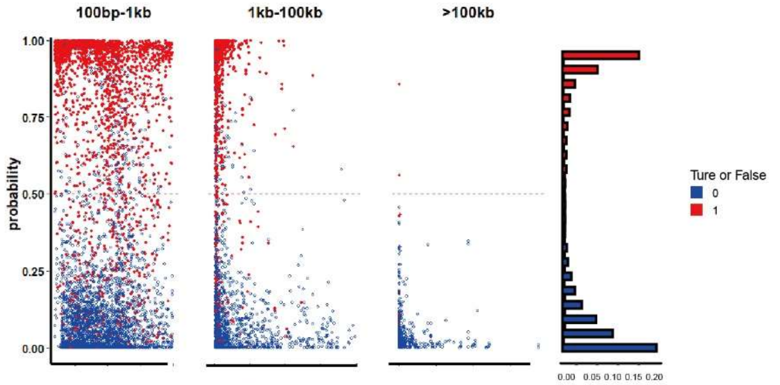


E) CNV-P_breakdancer

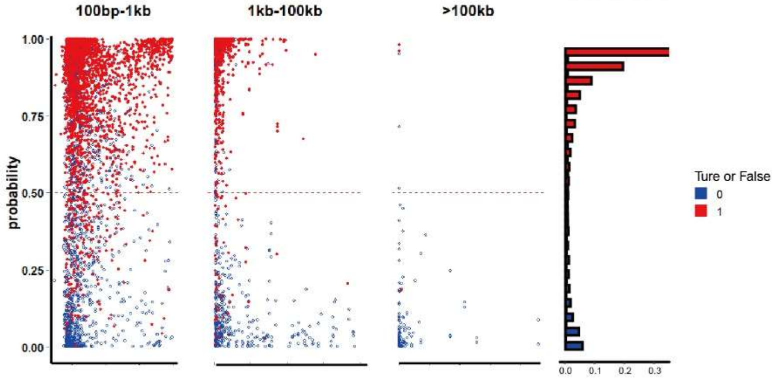

**Figure S6. Probability of CNV-P predictions at different size ranges.** The distribution of all CNV calls based on the probability scores, predicted by CNV-P across three size ranges is shown on left, and the frequency of validated and invalidated CNVs, distributed across 20 bins is shown on right. A) Lumpy; B) Manta; C)Pindel; D)Delly; E)breakdancer

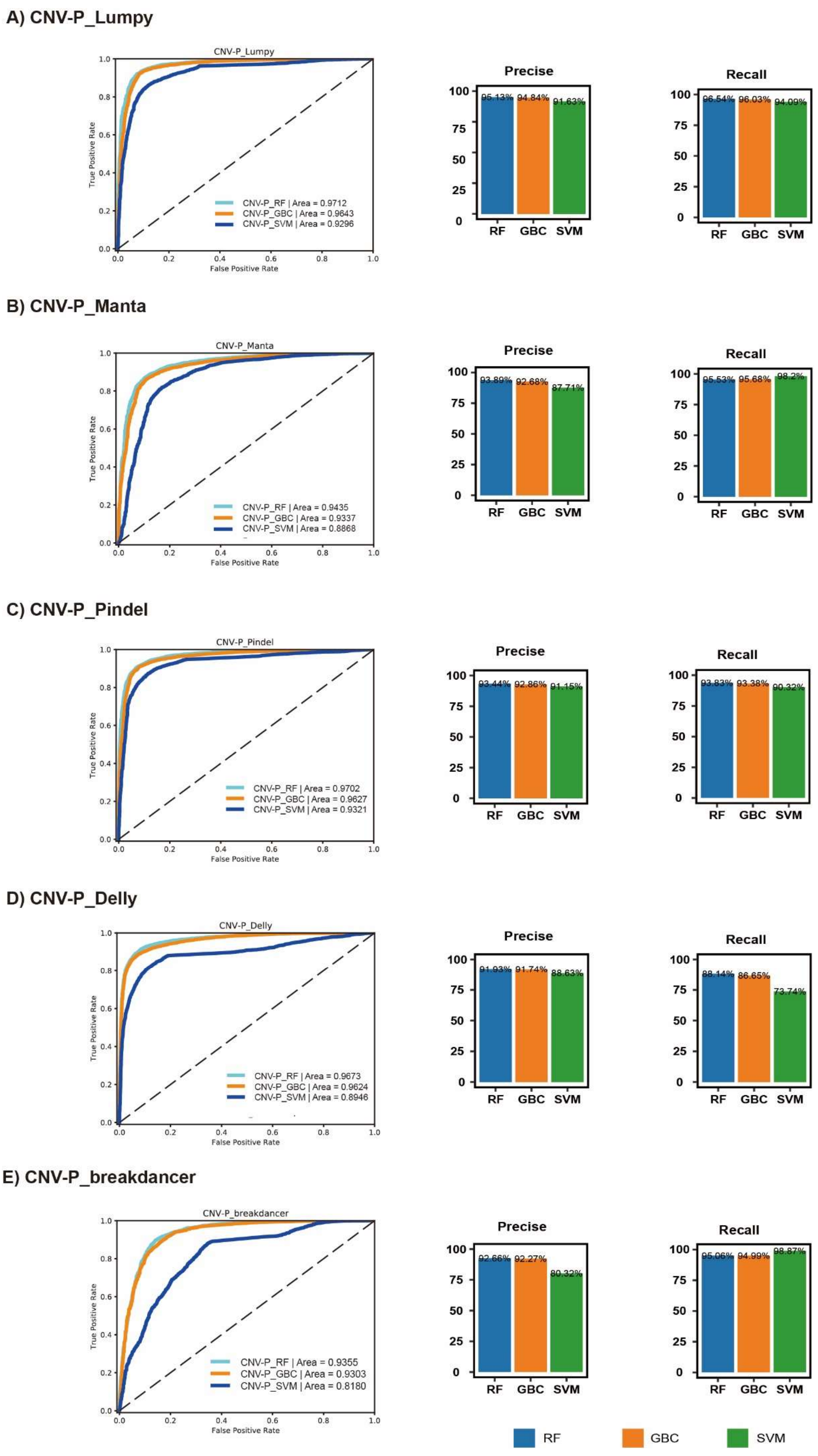
A) CNV-P_Lumpy
CNV-P_Lumpy
CNV-P_RF | Area = 0.9712
CNV-P_GBC | Area = 0.9643
CNV-P_SVM | Area = 0.9296
True Positive Rate
False Positive Rate
Precise
95.13% 94.84% 91.63%
Recall
96.54% 96.03% 94.09%
RF GBC SVM
B) CNV-P_Manta
CNV-P_Manta
CNV-P_RF | Area = 0.9435
CNV-P_GBC | Area = 0.9337
CNV-P_SVM | Area = 0.8868
Precise
93.89% 92.68% 87.71%
Recall
95.53% 95.68% 98.2%
C) CNV-P_Pindel
CNV-P_Pindel
CNV-P_RF | Area = 0.9702
CNV-P_GBC | Area = 0.9627
CNV-P_SVM | Area = 0.9321
Precise
93.44% 92.86% 91.15%
Recall
93.83% 93.38% 90.32%
D) CNV-P_Delly
CNV-P_Delly
CNV-P_RF | Area = 0.9673
CNV-P_GBC | Area = 0.9624
CNV-P_SVM | Area = 0.8946
Precise
91.93% 91.74% 88.63%
Recall
88.14% 86.65% 73.74%
E) CNV-P_breakdancer
CNV-P_breakdancer
CNV-P_RF | Area = 0.9355
CNV-P_GBC | Area = 0.9303
CNV-P_SVM | Area = 0.8180
Precise
92.66% 92.27% 80.32%
Recall
95.06% 94.99% 98.87%
RF GBC SVM

**Figure S7. Performance comparison of three classifiers supported by CNV-P.** For each CNV-P classifier A) Lumpy; B) Manta; C) Pindel; D) Delly; E) breakdancer. we illustrated ROC, precision values and recall values generated by RF, GBC and SVM. The results demonstrate the superior performance of the Random Forest-based classifier.

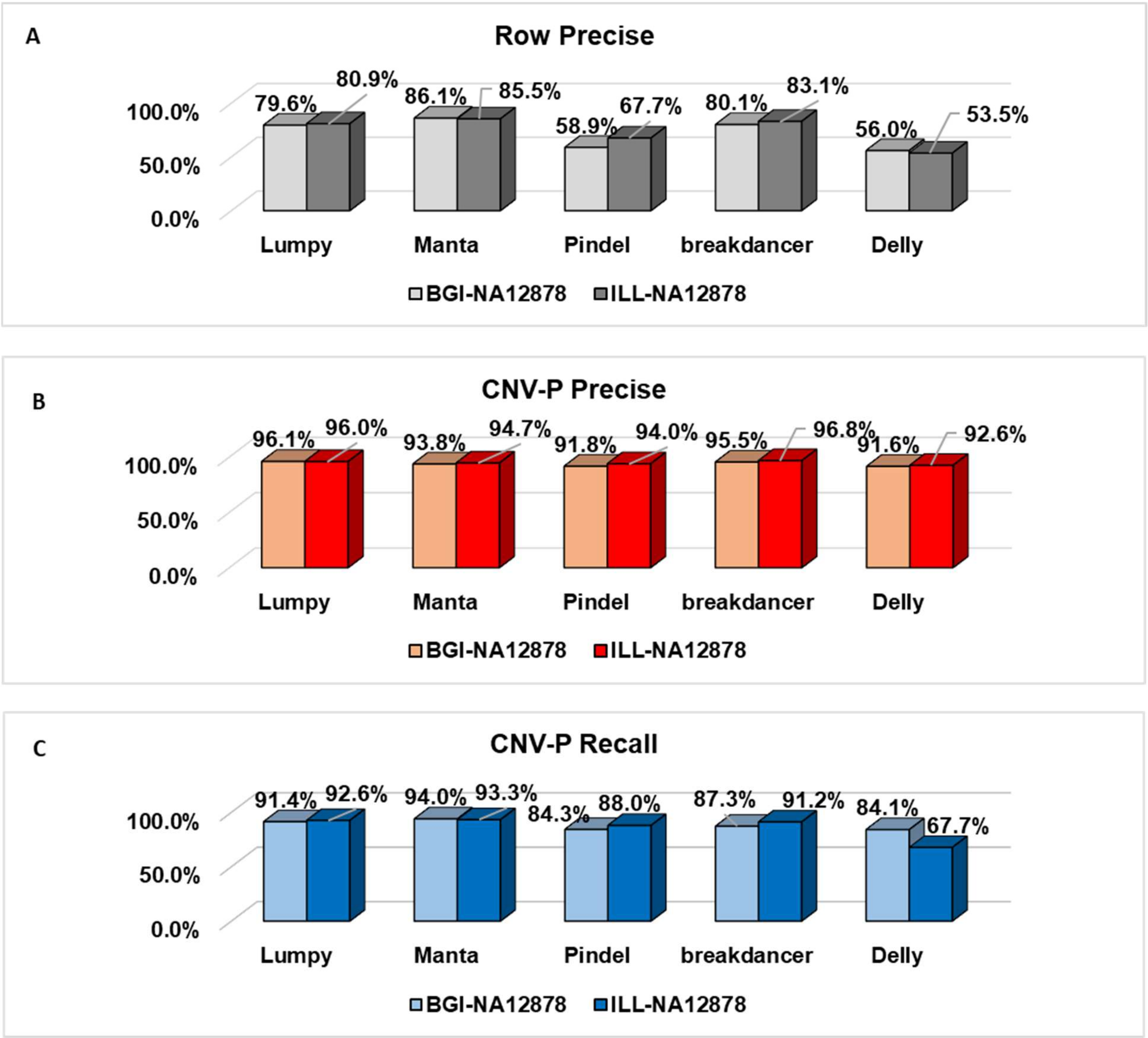


**Figure S8. CNV-P is robust across different sequencing platform.** The A)Row precise; B)CNV-P precise; C)CNV-P recall; across BGI-500 and Illumina sequencing platform.

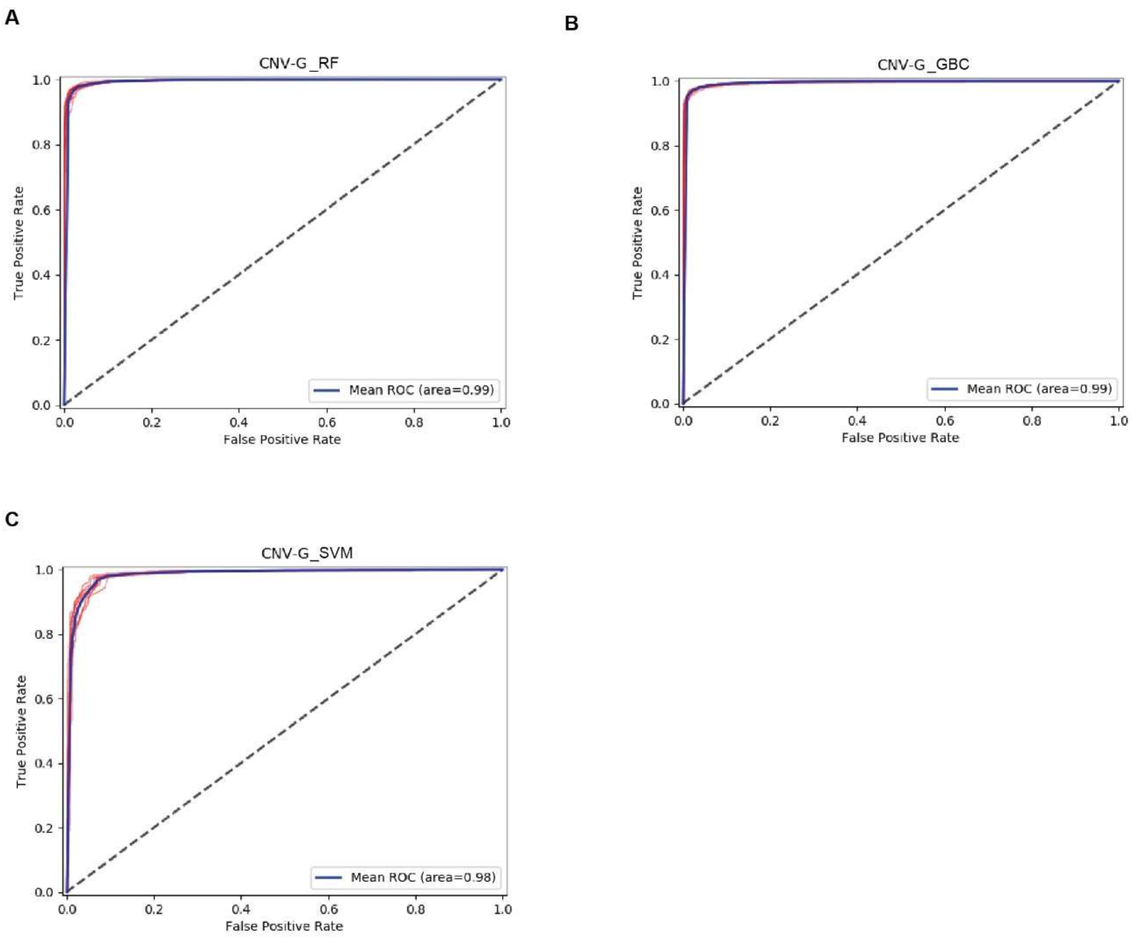


**Figure S9. 10-fold Cross validation for CNV-G.** The ROC of 10-fold Cross validation for A) RF-based; B) GBC-based; C) SVM-based CNV-G.
